# An accurate aging clock developed from the largest dataset of microbial and human gene expression reveals molecular mechanisms of aging

**DOI:** 10.1101/2020.09.17.301887

**Authors:** Vishakh Gopu, Ying Cai, Subha Krishnan, Sathyapriya Rajagopal, Francine R. Camacho, Ryan Toma, Pedro J. Torres, Momchilo Vuyisich, Ally Perlina, Guruduth Banavar, Hal Tily

## Abstract

Accurate measurement of the biological markers of the aging process could provide an “aging clock” measuring predicted longevity and allow for the quantification of the effects of specific lifestyle choices on healthy aging. Using modern machine learning techniques, we demonstrate that chronological age can be predicted accurately from (a) the expression level of human genes in capillary blood, and (b) the expression level of microbial genes in stool samples. The latter uses the largest existing metatranscriptomic dataset, stool samples from 90,303 individuals, and is the highest-performing gut microbiome-based aging model reported to date. Our analysis suggests associations between biological age and lifestyle/health factors, e.g., people on a paleo diet or with IBS tend to be biologically older, and people on a vegetarian diet tend to be biologically younger. We delineate the key pathways of systems-level biological decline based on the age-specific features of our model; targeting these mechanisms can aid in development of new anti-aging therapeutic strategies.

## Introduction

*Biological age* refers to biological markers of the aging process, and may be accelerated or slowed in some individuals relative to their chronological age. Recent research has proposed computational aging clocks based on various biomarkers including metabolites, blood cell count and other routine lab tests (Earls et al., 2019; Momoshina, Kochetov et al. 2018), DNA methylation (Fraga & Esteller, 2007; Horvath & Raj 2018; Bell et al. 2019), gene expression in tissue (Momoshina, Volosnikova et al. 2018) or blood (Harries et al., 2011; Lin et al., 2019), taxonomic composition of the gut microbiome (Galkin et al., 2020), among others. Aging clocks propose to use a signal derived from these biomarkers as a health-related metric for aging. In this paper we present two biological age metrics, one derived from the metatranscriptome of the gut microbiome, and one from the transcriptome of capillary blood. These two metrics together arguably capture the most comprehensive view of human biology.

Molecular markers from both microbial and human cells have been used to develop aging clocks. The composition and function of the gut microbiome changes with age, and may modulate healthy aging through multiple mechanisms. The increased dysbiosis associated with age can lead to innate proinflammatory immune responses, and the small molecules secreted by the gut microbiome affect host metabolism and signalling pathways that vary with age (see review in Kim & Jazwinski, 2018, i.a.). There is evidence that these microbiome changes over time are directly implicated in human healthspan. Maffei et al. (2017) show that certain properties of the gut microbiome, notably taxonomic diversity, are more predictive of a frailty index measuring mortality risk than is chronological age. Similarly on the human side, several molecular markers may modulate healthy aging. Perhaps the strongest aging clocks proposed so far have relied on biomarkers related to the epigenome such as DNA methylation. While these can act as an estimator of the biological age, they are not comprehensive and they have limited ability to pinpoint the regulators of the biological clock. Both the gut microbiome and the human molecular mechanisms are known to participate in widespread epigenetic interactions (see survey in Watson & Søreide, 2017), so a biological clock based on both of these functions can potentially inform specific therapeutic avenues to slow down aging. These may include personalized diets, supplements (vitamins, minerals, prebiotics, probiotics, food extracts, etc.), pharmaceuticals, phages, immunotherapies (vaccines, antibodies), etc.

While there are many ways to define biological age and operationalize the development of an aging clock (Jia et al., 2017), a common approach is to fit a machine-learned model to predict the chronological age of the human subject from the biomarker. This model’s predictions will deviate from chronological age to some extent: for example, if the subject’s biomarker profile is more similar to biomarker profiles of older people than to their peers, the model will overpredict age. The model’s predictions can be interpreted as a biological age in the sense that they approximate the age of a typical subject with the given biomarker profile. In this paper we show that the gut microbiome metatranscriptome as well as the blood transcriptome display strong associations with age to allow the creation of an aging clock.

## Methods

### Study cohorts^1^

Table 1 gives an overview of the stool microbiome and blood transcriptome cohorts used in this study.

**Table 1:** Model performance by cohort.

|  | Stool microbiome |  |  |  | Human blood transcriptome |
| --- | --- | --- | --- | --- | --- |
|  | Microbiome discovery cohort (cross-validated) | Microbiome Validation cohort (prospective) | Matched cohort with CV in Galkin et al. | Matched cohort with HC in Galkin et al. | Blood transcriptome discovery cohort (cross-validated) |
| Cohort size | 78,637 | 11,666 | 1164 | 252 | 1494 |
| Sex (% female) | 64.10 | 65.37 | 64.95 | 48.41 | 61.29 |
| Age (y) mean ± s.d. | 46.79 ± 15.9 | 43.22 ± 15.75 | 49.00 ± 15.35 | 48.06 ± 11.36 | 47.24 ± 14.22 |
| Baseline MAE | 12.98 | 12.90 | 13.03 | 9.36 | 11.75 |
| Prediction $R^2$ | 0.42 ± 0.00 | 0.46 | 0.46 | 0.31 | 0.53 ± 0.02 |
| Prediction MAE | 9.49 ± 0.02 | 9.21 | 9.14 | 7.64 | 7.63 ± 0.25 |

- The stool microbiome cohorts consist of samples obtained from unique customers of Viome’s Gut Intelligence product. These samples were divided into a discovery cohort of 78,637 samples, and a validation cohort of 11,666 samples.
- The Galkin et al. matched microbiome cohorts are intended to allow comparison of this model to the one presented in that work, and were constructed by randomly choosing one Viome customer from our validation set with the same age as each person in the Galkin et al. datasets. One person in the matched CV cohort could not be paired with a unique sample in our data, so our cohort has 1164 samples rather than Galkin et al.’s 1165. For the HC cohort we additionally matched on sex, which was impossible in the larger CV cohort.
- The human blood transcriptome cohort consists of samples obtained from 1494 unique customers of Viome’s Health intelligence product and associated research studies.

### Sample processing and bioinformatics

Stool samples were collected, preserved and processed using the metatranscriptomic method described in Hatch (2019). Paired-end reads were mapped to genomes (Breitwieser et al., 2017) and to a catalog of microbial genes with KEGG ortholog (KO) annotations (Kanehisa & Goto, 2000), and quantified using the expectation-maximization algorithm (Dempster, et al. 1977). This yields two views of the relative activity of each gut microbiome sample, one taxonomic and one functional. The taxonomic view aggregates reads to the species level, while the functional view aggregates the same reads to KOs.

Blood samples were collected, preserved and processed using the whole blood transcriptomic method described in Toma et al. This method is selective for polyadenylated RNA. Paired-end reads were mapped to the human genome. Gene expression levels were computed by aggregating transcripts per million estimates per gene using an approach based on Salmon version 1.1.0 (Patro et al., 2017), as described in (Toma et al. 2020).

### Machine learning

The microbiome data was transformed using the centered log ratio transformation (CLR) (Aitchison, 1986) after imputation of zero values using multiplicative replacement (Martín-Fernández et al., 2003). The human gene expression data was transformed using a Yeo-Johnson power transformation.

Machine-learned models for both stool microbiome and blood transcriptome are Elastic Nets (EN: linear regression with tunable L1 and L2 regularization). We also tried other approaches including deep neural networks (DNN), random forest, Adaboost and a combination of metric learning and k-nearest neighbors. As results were similar across all approaches, we report the simplest and most interpretable model class.

For the stool microbiome model, hyperparameter optimization was done using a 5-fold cross-validation on the discovery cohort. A final model using the optimal hyperparameter setting was then trained on the full discovery cohort and applied to the validation cohort to test generalization. Hyperparameter settings were scored using *R*^*2*^ and the selected model was evaluated using Mean Absolute Error (MAE) and *R*^*2*^.

For the human blood transcriptome model, model evaluation was performed using a nested CV due to the smaller dataset size. An inner 3-fold CV was used for hyperparameter selection while an outer 5-fold CV was used for model evaluation. Hyperparameter settings in each were scored using *R*^*2*^ and evaluated using MAE and *R*^*2*^.

Baseline MAE was computed as MAE from the median age of the cohort. The discovery cohort was used to tune model hyperparameters using cross validation, then a final model was trained on the full discovery cohort. The final model was applied to the validation cohort (drawn from the same population but not used in training), cohorts matched to two validation datasets (CV and HC) analyzed in Galkin et al. (2020), and additional cohorts described in Cohort Comparisons below.

### Cohort comparisons

As an additional exploration of our stool microbiome model, we compare the biological age predicted for a number of specific subsets of the full microbiome dataset (discovery plus validation) corresponding to populations of interest. In each case we select all available samples from the population of interest (e.g. vegetarians), and create an appropriate control cohort (e.g. omnivores) where each member of the control cohort is matched on age to one member of the reference cohort. We perform a paired sample t-test to determine whether there is a significant difference in biological age between the cohorts. The cohorts consist of: people reporting the special diets *vegan, vegetarian, organic, paleo, ketogenic* (contrasted with people following no special diet); people with self-reported *IBS* and *Diabetes* (contrasted with people reporting no health issues); and heavy *drinkers* (contrasted with non-drinkers), where heavy drinkers were defined following Mayo Clinic guidelines as consuming 15 or more drinks per week for males and 8 or more for females.

### Viome Functional Categories (VFCs)

We have built an annotation system that integrates both species level taxonomic activity and the functional expression profiles from KOs into higher order biological themes, called ‘Viome Functional Categories (VFC)’. VFCs are expert-curated themes that account for pathway directionality of feature association (activation /suppression, production/degradation, protective/harmful) and provide mechanistic insights into aging. Microbial taxonomic and gene expression features are grouped into 11 biological themes covering 32 VFCs. Human gene expression features are grouped into 26 themes covering 65 VFCs. For example, the theme “ProInflammatory Activities in Aging” consists of 8 different VFCs. Within this theme, the “Ammonia Production Pathways” VFC contains 3 KOs, all with a positive association with aging in the model. More details are provided in the Supplementary Materials.

## Results

Figure 1 presents descriptive statistics of the discovery cohort. Ages of sample donors range from <1 years to 104 years, with 2686 donors below 18 years of age (included with parental consent). Study participants come from over 60 countries (86% US, 8% Canada, 3% Australia, 1.5% EU, 1% UK, rest from other countries). We do not observe any differences in taxonomic richness by age (Fig 1b-c); nor do we find differences in taxonomic diversity or active function richness. None of these four measures were found to increase predictive accuracy when included in our models. Fig 1d-e shows the taxa at the species level and KOs that vary the most with age. To identify these, we calculated the mean CLR for each feature in each decade of age in 70% of the discovery cohort, and chose those with the highest variance across ages. Then we plotted the trend in mean CLR by decade in the remaining 30% of the data, grouped by Viome Functional Category (VFC). Notably, all of the KOs with the highest positive association with age are part of Methanogenesis Pathways resulting in production of methane gas.

**Figure 1.**
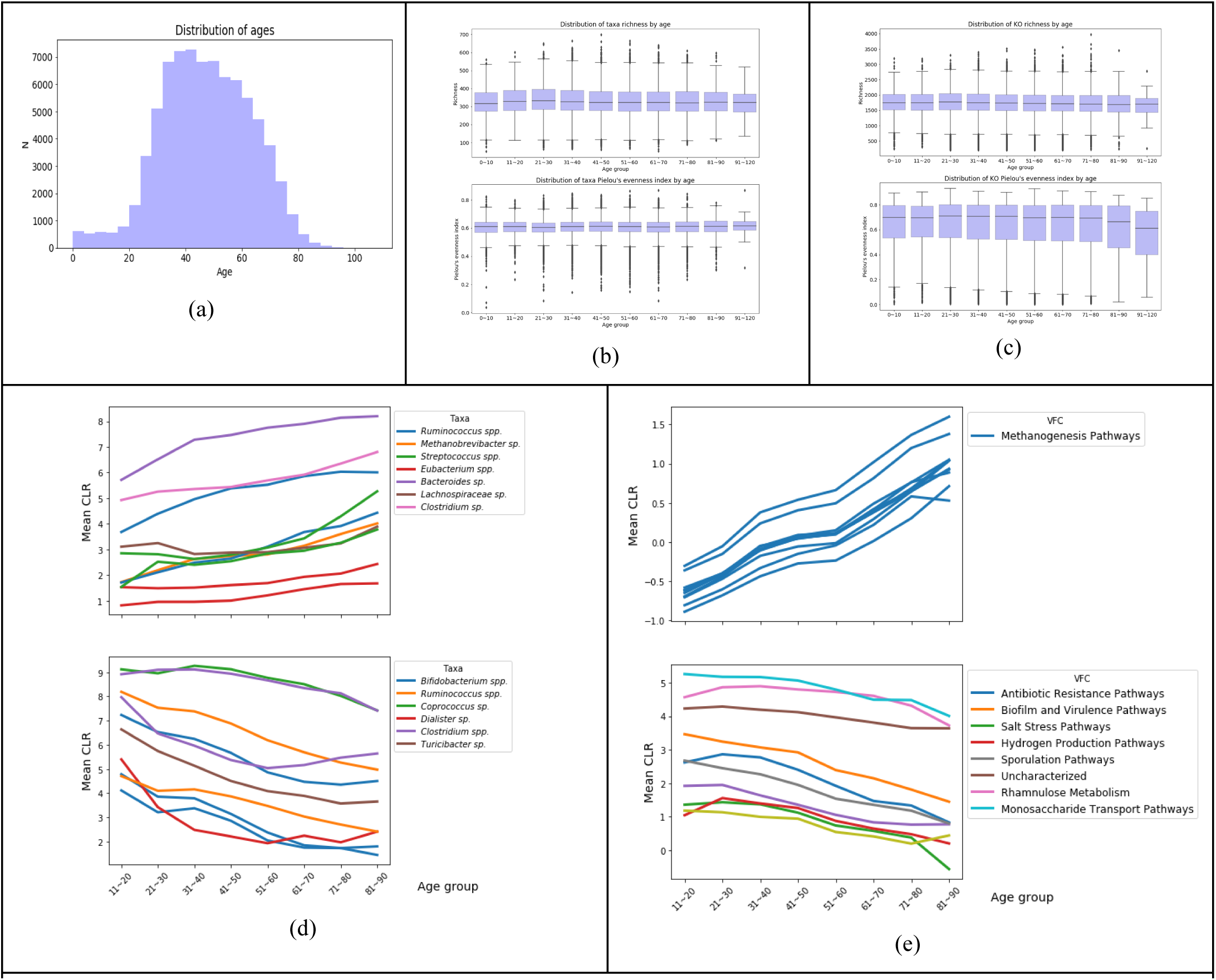
Descriptive statistics for the microbiome discovery cohort of Table 1. (a) age distribution (b) richness and shannon diversity of active microbial richness by decade (c) richness and Pielou’s evenness index for active functions (d-e) mean CLR transformed expression levels of species/KOs by age for most variable species/KOs grouped by genus/VFC.

Our biological age model’s performance is presented in Table 1. For the independent validation cohort, the model predicts chronological age above the baseline MAE of the datasets, and accounts for around 46% of the variance in age by *R*^*2*^, the standard metric of quality of fit in regression tasks. Our biological age model’s predictions and most important predictors are shown in Figure 2. Figure 2b shows the features with highest absolute coefficients (above 0.3 for taxa and 0.15 for KOs) grouped by Viome Functional Categories (VFCs) and further grouped into themes (see Supplementary Materials).

**Figure 2.**
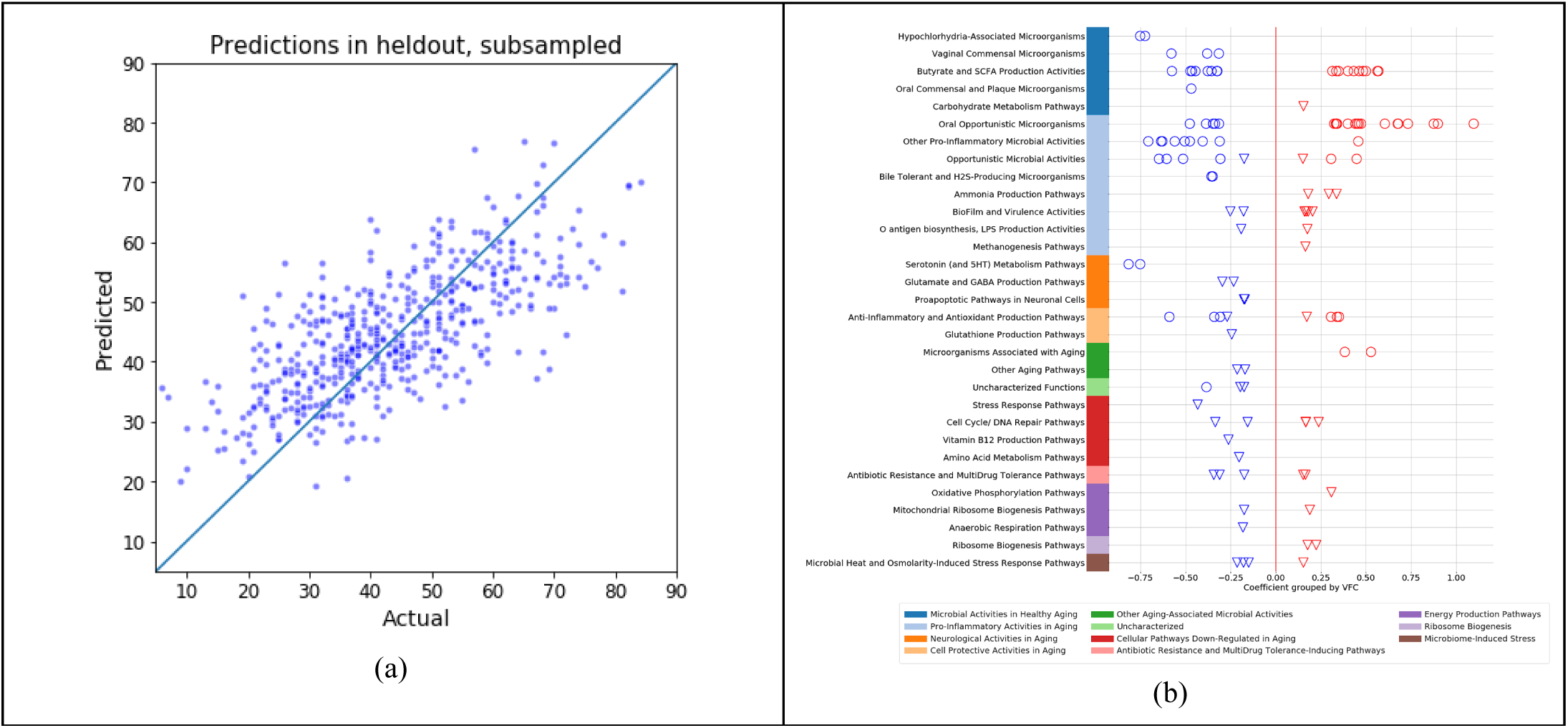
Biological aging model using the microbiome discovery cohort of Table 1. (a) Predicted vs. actual age in held-out validation data (for clarity, only a random subset of points is shown) (b) Coefficients for the microbial taxonomic features (circles) and KO features (triangles) grouped into curated Viome Functional Categories (VFCs)

Table 1 presents performance of the model under 5-fold cross validation. The model accounts for around 53% of variance in ages in the dataset. Figure 3 presents the predictions of our aging model based on human blood transcriptome.

**Figure 3.**
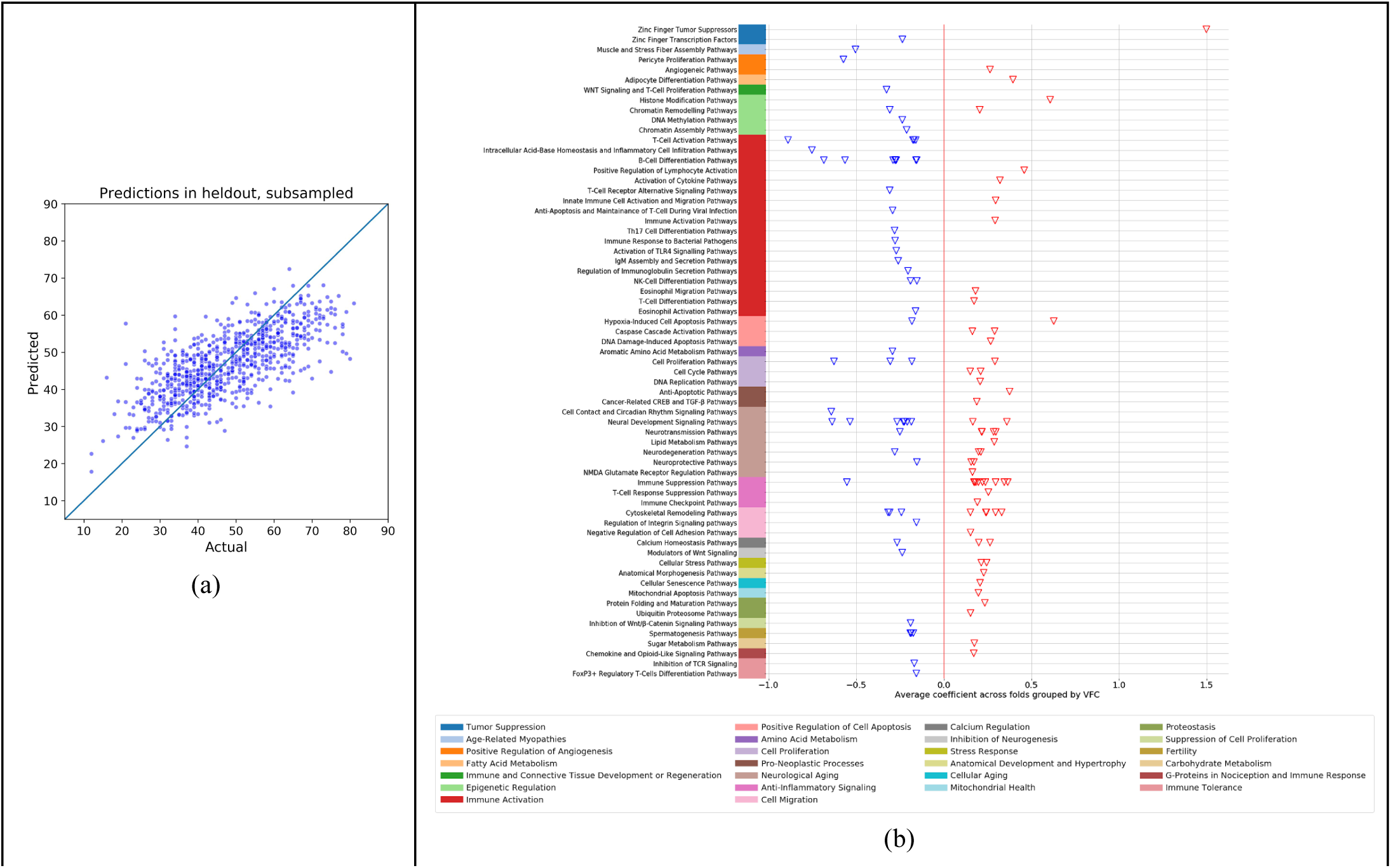
Biological aging model using the human blood transcriptome discovery cohort of Table 1. **(a)** Actual vs. Predicted **(b)** Top coefficients grouped by Viome Functional Categories (VFCs)

### Cohort comparisons

To explore the biological age of specific populations of interest, we present summaries of paired sample t-tests in Figure 4a, which depicts the difference in mean biological ages for specific cohort comparisons of interest, together with p-values from corresponding t-tests. Figure 4b shows the difference in chronological age for these same populations within the discovery cohort. We note that the model picks up several interesting differences between these populations and their age-matched controls. Vegetarians and vegans both tend to have a lower biological age than omnivores, while those following the ketogenic or paleo diets are biologically older than omnivores. Heavy drinkers are biologically older than non-drinkers. People with diabetes or IBS appear older than healthy controls. Some of these results may reflect chance patterns in the training data, as discussed below.

**Figure 4.**
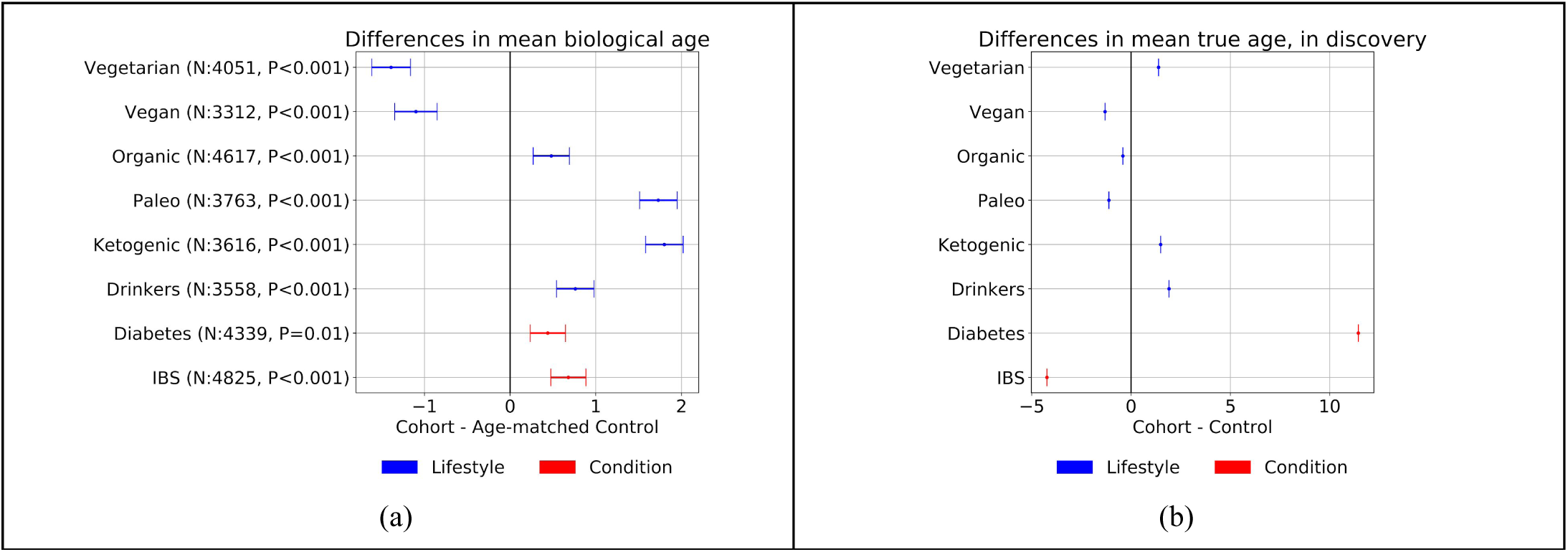
Cohort comparisons. (a) Mean and standard error of biological age differences between cohorts and age-matched controls where *p*-values < 0.1 from paired t-tests. (b) Mean and standard error of chronological age differences between cohorts and controls in the discovery cohort

## Discussion

### Model performance

The models presented here are capable of predicting chronological age above the baseline MAE of the datasets, and account for around 46% (stool) and 53% (blood) of the variance in age by *R*^*2*^. We note that some discrepancy between predicted and actual age is expected in a useful biological age candidate. If age was perfectly predicted, it would indicate either that the aspects of health captured by the biomarker decline in lockstep with chronological age, or that the biomarker is statistically associated with properties that vary systematically with age but are irrelevant to health.

We present an in-depth comparison of our microbiome model work with the gut metagenomic aging clock reported by Galkin et al. (2020). Galkin et al. report MAE of 10.60 and *R*^*2*^ of 0.21 (vs our 9.49 and .42) in a dataset with a baseline MAE of 13.03 (vs our 12.98). In a secondary validation exercise, their model obtains MAE of 6.81 and *R*^*2*^ of 0.134 when applied to a separate dataset (HC) with a lower baseline MAE of 9.27^2^. Since metatranscriptomic data is unavailable for that cohort, we created an additional validation cohort with exactly the same age distribution shown in Table 1. In this cohort, our model attains MAE of 7.64 (vs their 6.81) and *R*^*2*^ of .31 (vs their .13). *R*^*2*^ is the standard metric used for quality of fit in a regression task, and these numbers suggest that our metatranscriptomic model provides a better overall fit to this distribution of ages.

Interestingly, Galkin et al. report that an Elastic Net model was unable to extract significant signal from the data, and achieved their best performance using a deep neural net (DNN). In contrast, we found similar performance across model types, including when using a neural network architecture modeled after the one they report. Using a linear model is advantageous in terms of interpretability and actionability: the influence of each biological feature on the model’s age prediction is transparent, making it straightforward to determine a set of candidate targets to act on.

Although we report separate models for the microbiome and human gene cases, these models could be combined to give a single prediction. One straightforward way to do this would be to weight the predictions of each by the models’ precision. In future work we hope to collect a large dataset with both stool microbiome and human gene expression data collected simultaneously for the same users, which will allow us to fit a single joint model.

The resolution of the data supplied by our clinical grade and fully-automated lab analysis method allows identification of microorganisms at the strain level, although for this analysis we aggregate data to the species level. This contrasts with 16S gene sequencing, which does not discriminate between species within most genera. This additional resolution appears to be important to capture age-related variation. In several cases, some species of a genus are associated with older age, and others associated with younger age (shown in Figure S1 of supplementary materials).

Contrary to much published literature (de la Cuesta-Zuluaga et al., 2019; Hopkins et al., 2002; Mariat et al., 2009; Koenig et al., 2011; Yatsunenko et al., 2012), chronological age was not associated with significant changes in alpha diversity (richness or evenness) of taxa or KOs across decades (Figure 3b and Figure 3c) despite significant changes in individual taxa and KOs over time (Figure 3d and Figure 3e). This difference may be due to our RNA-based approach, whereas previous studies have used DNA-based approaches (amplicon or metagenomics). It is intriguing that the richness of active gene expression captured in this RNA-based data remains steady throughout life. It is possible that changes in taxonomic diversity observed in DNA-based approaches might help to retain a certain level of functional stability (Kang et al., 2015) obtained early in life.

### Cohort comparisons

Figure 4a shows that populations following certain lifestyle choices or suffering from specific conditions are systematically assigned a different biological age from their age-matched controls. For instance, those on plant-based diets appear younger (see review in Medawar et al., 2019). On the other hand, heavy alcohol drinkers and IBS and diabetes sufferers appear older. Those on a ketogenic or paleo diet tend to have a higher biological age than controls. For some of these cohorts (e.g. drinking, diabetes) the age difference in the training data has the same sign as the predicted age (Figure 4b). In these cases it is unclear whether the model has identified aspects of the microbiome that reflect general aging that are modulated in the groups of interest. However for other cohorts (i.e. vegetarians, paleo, IBS) the opposite pattern is seen, which shows that the microbiome features associated with these populations include features associated with aging in the general population. Overall, these results are consistent with an interpretation of our biological age metric as reflecting an accumulation of lifestyle choices and disease status that contribute to healthy aging.

### Molecular feature interpretation

The VFC are expert curated functional themes that provide mechanistic insights into the aging process from the features predictive of the biological age model. For instance, the Pro-Inflammatory Activities in Aging theme captures several VFCs like Oral Pathobionts, Ammonia Production Pathways and Methanogenesis Pathways that could lead to increased systemic inflammation and aging. The presence and activity of oral taxa in the gut is an established marker of hypochlorhydria (low stomach acid), which is known to develop with age as stomach acid levels and digestive efficiency progressively decline (Martinsen et al., 2005; Banoo et al., 2016; D’Souza, 2007). One of the common outcomes of low stomach acid is small intestinal bacterial overgrowth (SIBO), which allows overproduction of gases by oral and opportunistic microbes that should not have passed through the stomach environment, causing bloating and discomfort. SIBO can have several forms, depending on which gases are being mainly generated. A methane-dominant SIBO is characterized by excessive methanogenesis, which is also a significant pathway theme increasing with age. Methane is produced by Archaea, which has been reported to increase with age either due to relative decrease of other commensals or due to other factors allowing more methanogens to thrive in the gut. The increased ammonia and methane production contribute to dysbiosis, dysmotility, and pro-inflammatory effects that can be disruptive to the gut lining (Ni et al 2017). It is worthy to highlight that both KO and Taxa features of our model collectively contribute to microbial Proinflammatory Activities in Aging. On the other hand, Cell Protective Activities in Aging theme captures Anti-Inflammatory and Antioxidant Production Pathways, such as Glutathione Production Pathways, which are known to be diminished with aging (Homma et al 2015). Other than microbial Proinflammatory Activities, the model also predicts an association of Human Inflammatory Pathways with aging. Our model predicts increased demands on activity of pathways involved in B-cell differentiation, T-cell proliferation, T-cell differentiation, Eosinophil Migration, Cytokine Secretion, which can be viewed as the human response to activation of TLR4 and other signaling pathways with aging from microbial or environmental origins. To mitigate proinflammatory challenges, activation of innate immune response must be a precise and well-timed interplay between various immune cells on a molecular level (Takatsu, 1997). While some of the proinflammatory activities increase with age, however, many of the crucial T-cell response elements actually decrease in expression, which can result in insufficient ability of the immune system to respond to the sources of inflammation with age.

The Microbial Activities in Healthy Aging theme captures VFCs like Vaginal Commensal Microbes, Butyrate and Short-Chain Fatty Acid (SCFA) Production and Oral Commensal and Plaque Microbes that the model finds to be negatively associated with aging. The Cellular Pathways Downregulated in Aging theme captures other pathways that decrease with aging like Vitamin B12 production and Amino acid metabolism that are important for maintaining the gut diversity and homeostasis. Increased gut dysbiosis mediated by senescence-associated inflammatory conditions also results in the decreased microbial metabolite production (Gargari et al 2018). Stress Response Pathways also generally decline and become less efficient with age, while sources of Cellular Stress increase. (Edwards, 2011; Calderwood, 2009).

The role of gut microbiome in neuro-generative processes is increasingly evident, and the perturbation of the microbiome and microbial products has been demonstrated to affect behavior (Hsaio et al 2013) through the well-known gut-brain axis (Dumitrescu et al 2018). In the Neuronal Activities in Aging theme, the VFCs Glutamate and Gamma Amino Butyric Acid (GABA) Pathways, Serotonin Metabolism Pathways and Pro-apoptotic Pathways in neuronal cells show association with aging from the model in line with current knowledge of Neuroinflammation and cognitive aging. The inhibitory motor function mediated by GABA has also shown to be gradually declining with advancing age (Pauwels et al 2018), and serotonin loss leads to behavioral changes commonly observed in the elderly population (Meltzer et al 1998).

Additional supporting evidence for the declined neurotransmission and neuronal development with aging comes from the model’s predictive human gene expression features, which play a role in Neurotransmission Pathways, Neurodegeneration Pathways, and inverse relationship with Neuronal Growth and Development with aging themes.

Collectively, there are several integrative themes suggested by the significant KOs, Taxa, and human gene expression features of the model revealing a mechanistic systems biology viewpoint on aging. Some of the features have already been shown to play a role in age-related decline. For example, even though there may be a proinflammatory mechanism triggering certain immune responses from the gut microbiome, various other factors suggest inefficient or imprecise immune responses that may be needed to mitigate the sources of inflammation due to underactive T-Cell attachment to target cells. T-Cell populations may not only decline with age, but have also been suggested to change functional as they acquire more senescent-like characteristics (Quinn et al 2018). The mechanisms of T-Cell functional changes with age together with senescence are well-represented in the features of our model that belong to these themes. Various pro-neoplastic and proinflammatory themes are accompanied by stress response, senescence, epigenetic changes, and diminished T-Cell and other lymphocyte functions, which imply insufficient ability to maintain cellular health and immune surveillance needed to mitigate and clear any pathogens, cancer cells, or proinflammatory debris with age. This, in turn, can contribute to neuronal proinflammatory processes, which, together with the microbiome component, can promote low-grade neuroinflammation, progressive damage, and subsequent cognitive decline with age.

Some prior work has attempted to validate the relevance of biological age metrics to health by relating them to measures of health or mortality (e.g. Holly et al., 2013; Earls et al., 2019), and we are investigating a similar approach in ongoing work. We note that biological age may in general serve as a proxy for longevity or healthspan. Additionally, our results uncover novel machine-learned associations between age and specific metatranscriptomic features that could guide the design of nutritional interventions to reduce biological age and increase the human healthspan (see e.g. Sae-Lee et al., 2018; Ghosh et al., 2019). We will continue to evaluate and improve the models as we obtain more data.

## Conclusions

A major contribution of this paper is the development and validation of two models for biological age. Our stool metatranscriptome model uses what is to date the largest published cohort of stool microbiome samples, 90,303 in total. Although our human transcriptome model is based on a smaller dataset, it performs very well. Both models are capable of predicting chronological age well above the baseline MAE of the datasets, and account for 46% and 53% of the variance in age by *R*^*2*^ respectively. Using this standard metric for quality of fit in a regression task, the performance of our stool metatranscriptomic aging model is the best of any published stool microbiome aging clock, and our blood transcriptome-based model is the best for any models developed using large (N>1000) datasets, to our knowledge.

Another contribution of this paper is the biological age characterization of populations following specific dietary and lifestyle choices. While not all of these initial results are readily interpretable, the trends in some populations indicate a relationship between health and biological age - for example, those on a vegetarian diet appear younger, while those on a paleo diet or suffering from IBS appear older. These results are consistent with an interpretation of our biological age metric as reflecting an accumulation of lifestyle choices and disease status, and suggest that the microbiome features associated with these populations include features associated with aging in the general population.

Moreover, pathway analysis of the features and interpretation of functional themes suggests mechanistic insights predictive of aging, and further connects age-related microbial activities with human cellular expression patterns on a molecular level. This predictive signature not only offers additional clues to the role of immune function in progressive systems-level decline, but also provides possibilities for future nutritional or pharmacological anti-aging interventions.

This is the first report of using functional (i.e. gene expression) microbiome features and combining them with human gene expression data to build an accurate aging model. These microbial and human gene expression features open up the possibility to slow down human aging with specific therapeutic modalities that include natural (diet, supplements, lifestyle) and pharmaceutical (small molecules, phages, probiotic engineering, immunotherapies) approaches.

## Supplementary materials

**Figure S1.**
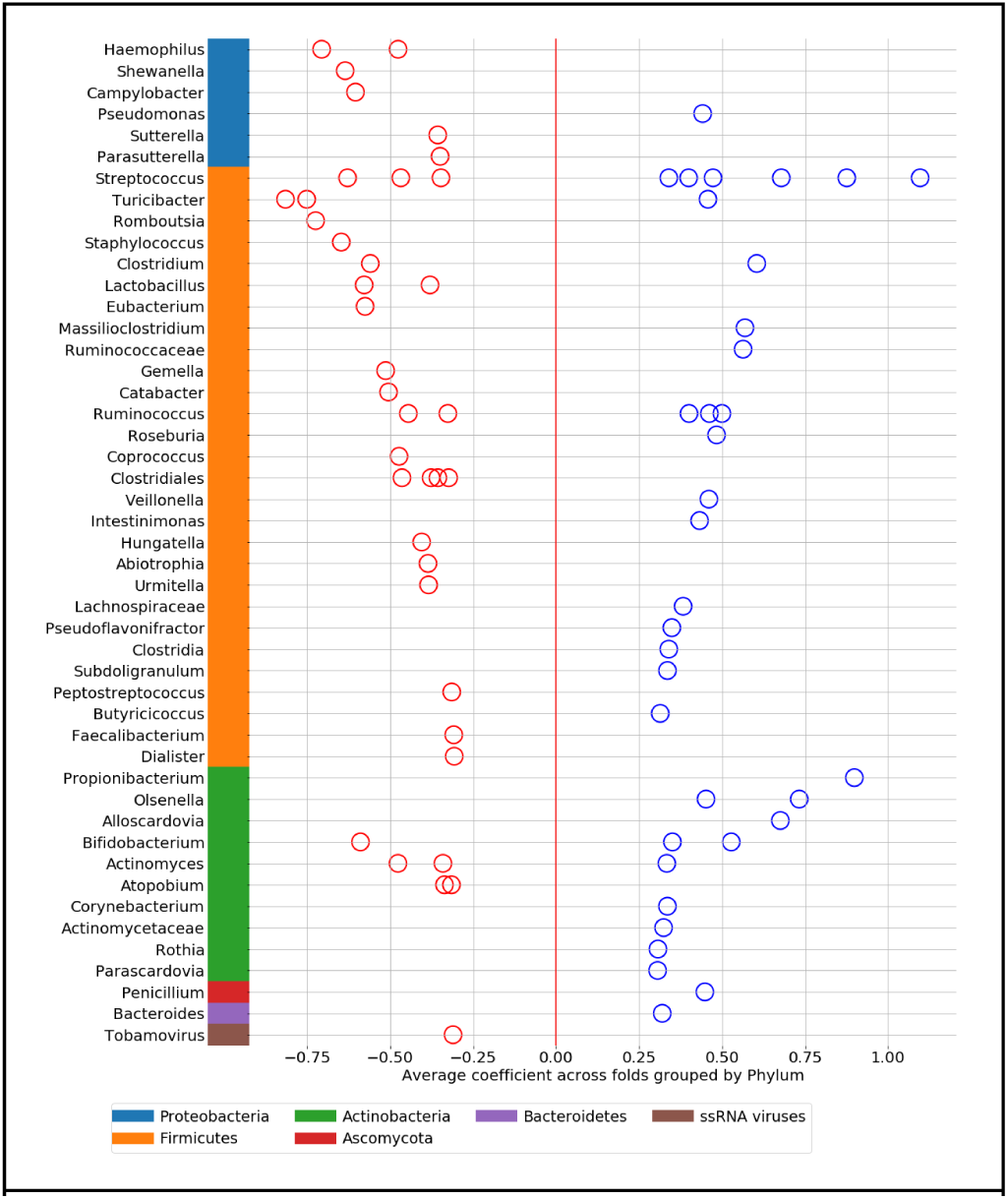
For the predictive model with microbiome discovery cohort, species features grouped into genus level, with absolute coefficients > 0.3

### Viome Functional Categories (VFC)

We have built an annotation system that integrates both taxonomic abundances and the functional expression profiles from KOs into higher order biological themes that are relevant for aging. A similar system has been built for human gene expression profiles. We call these biological themes Viome Functional Categories (VFC). The VFCs are highly curated themes that attempt to provide mechanistic insights into the aging process from the features predictive of the biological age model. We report a total of 32 VFCs grouped into 11 broad biological themes. The VFCs and the themes are as discussed below.

#### 1. ProInflammatory Activities in Aging

The ProInflammatory Activities in Aging theme provides evidence of a modified microbial synergy and dysbiosis of the gut bacterial involvement in the aging process. The term “Inflamm-aging” has been used to describe the proinflammatory environment that exists in older adults characterized by high concentrations of proinflammatory cytokines and impaired immune responses to pathogens and vaccines (Franceschi et al 2000). This theme captures some of the microbial contributions to aging related inflammation. Proinflammatory and opportunistic microbes such as Streptococcus sp., Propionibacterium sp., Rothia sp. and their associated functional components involved in increased virulence, such as the iron transport protein A (Carton et al 2006) and fatty acid double bond hydratase (Volkov et al 2010) may contribute to process of inflammation and aging (Nagata et al 2000, Maraki et al 2014).

Studies have reported an increased muscle and blood ammonia content with age, implicating the possibility of elevated purine nucleotide deamination during senescence (Mohan et al 1987). In addition, the microbial ammonia production pathway could contribute to increased ammonia levels in the aged population thereby increasing inflammation. Three KOs involved in ammonia production are predictive of age in the model.

#### 2. Antibiotic Resistance and MultiDrug Tolerance Inducing Pathways

Infectious diseases remain a serious problem for older people and the frequent prescribing of antibiotics led to the emergence of highly resistant pathogens among geriatric patients, such as methicillin-resistant Staphylococcus aureus, penicillin-resistant Streptococcus pneumoniae, vancomycin-resistant enterococci, and multiple-drug-resistant gram-negative bacilli (Yoshikawa 2002). Two antibiotic resistance and multidrug tolerance associated KOs are found to be associated with increased age.

#### 3. Microbial Pathways Down regulated in Aging

One KO each from Stress Response Pathway (HSP20), Vitamin B12 Production Pathway, Hydroxyproline and Proline Catabolism Pathway and two KOs from Cell cycle and DNA Repair Pathways are found to be associated with younger age. Three other KOs from Cell cycle and DNA Repair Pathways show an association with older age.

#### 4. Cell Protective Activities in Aging

The enzymatic and non-enzymatic cellular antioxidants such as superoxide dismutases, catalases, glutathione system, thioredoxin system, peroxidase systems, flavohemoglobins and nitrate or nitrite reductases coordinate the balance between the production and degradation of Reactive Oxygen Species (ROS) and Reactive Nitrogen Species (RNS) (Staerck et al 2017). Decreased GSH levels are associated with aging as well as in neurodegenerative disorders (Homma et al 2015). The model identifies a KO belonging to the GSH production pathway to be predictive of younger age. Two other KOs involved in anti-inflammatory and antioxidant functions (such as peroxiredoxin and superoxide dismutase) are also predictive of younger age.

#### 5. Microbial Activities in Healthy Aging

It is well known that gut commensals have significant influence on the host nutrition and metabolism by producing microbial metabolites like SCFAs, vitamins and cofactors that are readily absorbed from the intestinal epithelium and are also essential for healthy gut lining. The preservation of host-microbes homeostasis can counteract inflammaging, intestinal permeability and decline in bone and cognitive health (Biagi et al 2016). These microbial metabolites contribute to healthy aging and human longevity by maintaining gut diversity and homeostasis. Increased gut dysbiosis mediated by senescence-associated inflammatory disorders results in the decreased microbial metabolite production (Gargari et al 2018). This phenomenon has been captured in the features from the predictive model where SCFA producers such as Clostridiales sp. and Ruminococcus sp., Coprococcus sp., and Eubacterium sp. are predictive of younger age. Two vaginal commensal Lactobacilli and Atopobium, the beneficial bacteria detected in the vaginal microbiome of reproductive age women (Ravel et al 2001) is found in the feature to be predictive of younger age. Thus, reduced microbiome-related metabolic capacity in old age, such as lower levels of short-chain fatty acids (SCFAs), may also be associated with aging-related maladies such as irregular bowel transit, weight loss, cognitive decline, hypertension, vitamin D deficiency, diabetes, arthritis (Zapata et al 2015).

However, it is interesting to see that Ruminococcaceae sp., Ruminococcus sp., Roseburia sp., Intestinimonas sp., Pseudoflavonifractor sp., Subdoligranulum sp., Butyricicoccus sp., some of the well-known butyrate and SCFA producers are also predictive of older age. The presence of Ruminococcus, Bifidobacteria and Lachnospiraceae has been shown to increase in the elderly and associated with aging (Biagi et al 2016, Claesson et al 2012, Zapata et al 2015).

#### 6. Neurological Activities in Aging

The role of gut microbiome in neuro-generative processes is increasingly evident and the perturbation of the microbiome and microbial products have been demonstrated to affect behavior (Hsaio et al 2013) through the gut-brain axis (Dumitrescu et al 2018). Studies have reported differences in short-chain fatty acids synthesis, tryptophan metabolism, and synthesis/degradation of neurotransmitters to be associated with neurological and social problems (Zhu et al 2020). Abnormalities of neurotransmitter systems, especially the aberrations of neuronal signaling involving dopamine, glutamate, and γ-aminobutyric acid (GABA) are among the pathways known to be dysregulated in diseases like Schizophrenia (Zhu et al 2020).

KO and taxa features involved in neurotransmitters and neuronal signaling are also predictive of old age. For instance, two KOs involved in the glutamate and GABA production have a negative association with aging. Along with neurotransmitters, KOs involved in neurosteroid production pathways in the brain and the brain lipid metabolism resulting in the accumulation of propionic acid have also been negatively associated with age. One Streptococcus sp. identified by the model has previously been implicated in social behavior deficits, by altering neurotransmitter levels in peripheral tissues in recipient mice (Zhu et al 2020). This taxon feature is found to be positively associated with age.

Similarly, based on the human gene expression, we report 65VFCs that could be grouped into 26 aging relevant functional themes. Among the model’s most predictive features are genes belonging to the following categories: Proinflammatory Pathways, Neurological Aging Pathways, Epigenetic Regulation Pathways, and Cell Apoptosis Pathways.

#### 7. Human Inflammatory Pathways and Aging

A balanced proinflammatory and anti-inflammatory mechanism is the key for good immune health. An increased production of proinflammatory cytokine production leading to chronic proinflammatory environment is characteristic of aging and Immunosenescence. IL-18 is a proinflammatory cytokine, known to enhance TH1 and TH2 differentiation and immune response via stat/IFNgamma pathway activation (Dinarello, C.A., 1998). PTGER2 upon activation significantly enhances inflammation by inducing the expression of proinflammatory factors, such as interleukin (IL)-1β and IL-6 in tumor cells (Sun X., 2018). Aging could also be characterized by the decline in immune responses (Fuentes E., 2017). Notably, our model identifies an association between older age and immune suppressors such as TCR pathway inhibitors and immune checkpoint markers (Manieri, N.A, 2016, Vazquez-Cintron,E.J., 2012, Monti-Rocha, R., 2019). Several signaling pathways which are involved in T-cell proliferation and maturation is also associated with aging phenotype (Lyons, G.E., 2006, Zhang, N. 2011).

#### 8. Epigenetic Regulation and Aging

The model identifies pathways involved in Epigenetic Modification (Pal, S., 2016, Sun, L., 2018). Significant changes in chromatin and DNA organization with aging have been previously reported (Pagiatakis, C.,2019). The model identifies an association between age and Chromatin Assembly Pathways, Chromatin Remodeling Pathways, and DNA Methylation Pathways.

#### 9. Neurological Aging Pathways and Aging

Aging is characterized by neurodegeneration and gradual decline in cognitive functions. Our model detects association of several pathways involved in neurological aging, such as Neuroprotective Pathways, Neurodegeneration Pathways, Neurotransmission Pathways with aging. Interestingly many of the predicted features have also been previously reported to be associated with Alzheimer’s phenotype (Duron E, 2012, Bossù P, 2007, Reitz C, 2013).

## Footnotes

1 All participants consented to participation, and the study protocol was approved by a federally-accredited Institutional Review Board (IRB). All data was de-identified for the purpose of the analyses reported here.

2 Galkin et al. also report MAE of 5.91 for this dataset but note that the data contains multiple samples from many of the subjects, and after merging duplicates into averaged samples, performance falls to 6.85. As performance on averaged samples is not representative of performance on individual samples, we report metrics for the Galkin et al. model after randomly excluding all but one sample from each donor. The metrics we report here are calculated from the predictions for individual samples shared as part of that paper’s supplementary data.

## References

Aitchison J. The Statistical Analysis of Compositional Data. (1986); Chapman and Hall.

Aas, Jørn A., Bruce J. Paster, Lauren N. Stokes, Ingar Olsen, and Floyd E. Dewhirst. Defining the Normal Bacterial Flora of the Oral Cavity. Journal of Clinical Microbiology 43, no. 11 (November 2005): 5721–32. doi: 10.1128/JCM.43.11.5721-5732.2005.

Banoo, H. and Nusrat, N. Implications of Low Stomach Acid: An Update RAMA Univ. J. Med Sci (2016); 2(2): 16–26

Bell, C.G., Lowe, R., Adams, P.D. et al. DNA methylation aging clocks: challenges and recommendations. Genome Biol (2019); 20, 249. doi: 10.1186/s13059-019-1824-y

Breitwieser, F.P., Lu, J., Salzberg, S.L., A review of methods and databases for metagenomic classification and assembly, Briefings in Bioinformatics, 20(4), 1125–1136, doi: 10.1093/bib/bbx120

Calderwood, S.K., Murshid, A., and Prince, T. The Shock of Aging: Molecular Chaperones and the Heat Shock Response in Longevity and Aging – A Mini-Review. Gerontology. 2009 Sep; 55(5): 550–558.

de la Cuesta-Zuluaga J., Kelley S.T., Chen Y., Escobar J.S., Mueller N.T., Ley R.E., McDonald D., Huang S., Swafford A.D., Knight R, Thackray V.G.. Age-and sex-dependent patterns of gut microbial diversity in human adults. mSystems. 2019;4:e00261–19. doi: 10.1128/mSystems.00261-19

Dumitrescu, Laura, Iulia Popescu-Olaru, Liviu Cozma, Delia Tulbă, Mihail Eugen Hinescu, Laura Cristina Ceafalan, Mihaela Gherghiceanu, and Bogdan Ovidiu Popescu. “Oxidative Stress and the Microbiota-Gut-Brain Axis.” Oxidative Medicine and Cellular Longevity 2018 (2018): 2406594. https://doi.org/10.1155/2018/2406594.

Dempster, A.P., N.M. Laird and D.B. Rubin. Maximum likelihood from incomplete data via the EM algorithm. Journal of the Royal Statistical Society. (1977); Series B. (29), 1–37.

D’Souza, A L. Aging and the Gut. Postgraduate Medical Journal 83, no. 975 (2007): 44–53. doi: 10.1136/pgmj.2006.049361.

Earls, J.C., Rappaport, N., Heath, L,, Wilmanski, T., Magis, A.T., Schork, N.J., Omenn, G.S., Lovejoy, J., Hood, L., Price, N.D., Multi-Omic Biological Age Estimation and Its Correlation With Wellness and Disease Phenotypes: A Longitudinal Study of 3,558 Individuals, The Journals of Gerontology. (2019); 52–S60, doi: 10.1093/gerona/glz220

Edwards, H.V., Cameron, R.T and Baillie, G.S. The Emerging Role of HSP20 as a Multifunctional Protective Agent. Cell Signal. (2011); 23(9), 1447–54.

Fraga, M.F., and Esteller, M. Epigenetics and aging: the targets and the marks. Trends Genet. (2007); 23, 413–418.

Galkin, F., Mamoshina, P., Aliper, A., Putin, E., Moskalev, V., Gladyshev, V.N., Zhavoronkov, A. Human gut microbiome aging clock based on taxonomic profiling and deep learning, ISCIENCE (2020); doi: 10.1016/j.isci.2020.101199.

Ghosh T.S., Rampelli S., Jeffery I.B., et al. Mediterranean diet intervention alters the gut microbiome in older people reducing frailty and improving health status: the NU-AGE 1-year dietary intervention across five European countries. Gut (2019); doi: 10.1136/gutjnl-2019-319654

Harries LW, Hernandez D, Henley W, et al. Human aging is characterized by focused changes in gene expression and deregulation of alternative splicing. Aging Cell. (2011); 10(5):868–878. doi: 10.1111/j.1474-9726.2011.00726.x

Holly A.C., Melzer D., Pilling L.C., et al. Towards a gene expression biomarker set for human biological age. Aging Cell. (2013); 12(2):324–326. doi: 10.1111/acel.12044

Homma, Takujiro, and Junichi Fujii. “Application of Glutathione as Anti-Oxidative and Anti-Aging Drugs.” Current Drug Metabolism 16, no. 7 (2015): 560–71. https://doi.org/10.2174/1389200216666151015114515.

Hsiao, Elaine Y., Sara W. McBride, Sophia Hsien, Gil Sharon, Embriette R. Hyde, Tyler McCue, Julian A. Codelli, et al. “Microbiota Modulate Behavioral and Physiological Abnormalities Associated with Neurodevelopmental Disorders.” Cell 155, no. 7 (December 19, 2013): 1451–63. https://doi.org/10.1016/j.cell.2013.11.024.

Hopkins M.J., Sharp R., Macfarlane G.T.. 2002. Variation in human intestinal microbiota with age. Dig Liver Dis 34(Suppl 2):S12–S18. doi: 10.1016/S1590-8658(02)80157-8.

Horvath, S., & Raj, K. DNA methylation-based biomarkers and the epigenetic clock theory of aging. Nature Reviews Genetics (2018); 19, 371–384.

Jia L, Zhang W, Chen X. Common methods of biological age estimation. Clin Interv Aging. (2017); 12:759–772. doi: 10.2147/CIA.S134921

Kanehisa, M., Goto, S., KEGG: Kyoto Encyclopedia of Genes and Genomes, Nucleic Acids Research. (2000); 28(1), 27–30, doi: 10.1093/nar/28.1.27

Kim S. & Jazwinski S.M. The gut microbiota and healthy aging: A mini-review. Gerontology (2018); 64:513–520. doi: 10.1159/000490615

Koenig J.E., Spor A., Scalfone N., Fricker A.D., Stombaugh J., Knight R., Angenent L.T., Ley R.E. 2011. Succession of microbial consortia in the developing infant gut microbiome. Proc Natl Acad Sci U S A 108(Suppl 1):4578–4585. doi: 10.1073/pnas.1000081107.

Lin H., Lunetta K.L., Zhao Q., et al. Whole Blood Gene Expression Associated With Clinical Biological Age. J Gerontol A Biol Sci Med Sci. (2019); 74(1):81–88. doi: 10.1093/gerona/gly164

Mariat D., Firmesse O., Levenez F., Guimarăes V., Sokol H., Doré J., Corthier G., Furet J-P. 2009. The Firmicutes/Bacteroidetes ratio of the human microbiota changes with age. BMC Microbiol 9:123. doi: 10.1186/1471-2180-9-123.

Martín-Fernández, J.A., Barceló-Vidal, C. & Pawlowsky-Glahn, V. Dealing with zeros and missing values in compositional data sets using nonparametric imputation. Mathematical Geology. (2003); 35, 253–278. doi: 10.1023/A:1023866030544

Maffei V.J., Kim S., Blanchard E. 4th, et al. Biological aging and the human gut microbiota. J Gerontol A Biol Sci Med Sci. (2017); 72(11):1474–1482. doi: 10.1093/gerona/glx042

Mamoshina P., Kochetov K., Putin E., et al. Population specific biomarkers of human aging: A big data study using South Korean, Canadian, and Eastern European patient populations. J Gerontol A Biol Sci Med Sci. (2018); 73(11): 1482–1490. doi: 10.1093/gerona/gly005

Mamoshina P., Volosnikova, M., Ozerov, I.V., Putin, E. Skibina, E., Cortese, F., Zhavoronkov, A. Machine learning on human muscle transcriptomic data for biomarker discovery and tissue-specific drug target identification. Front. Genet. (2018); 9: 242

Martinsen, Tom C., Kåre Bergh, and Helge L. Waldum. Gastric Juice: A Barrier Against Infectious Diseases. Basic & Clinical Pharmacology & Toxicology 96, no. 2 (2005): 94–102. doi: 10.1111/j.1742-7843.2005.pto960202.x.

Medawar, E., Huhn, S., Villringer, A. et al. The effects of plant-based diets on the body and the brain: a systematic review. Transl Psychiatry 9, 226 (2019). doi:10.1038/s41398-019-0552-0

Meltzer, Carolyn Cidis, Gwenn Smith, Steven T. DeKosky, Bruce G. Pollock, Chester A. Mathis, Robert Y. Moore, David J. Kupfer, and Charles F. Reynolds. “Serotonin in Aging, Late-Life Depression, and Alzheimer’s Disease: The Emerging Role of Functional Imaging.” Neuropsychopharmacology 18, no. 6 (June 1998): 407–30. https://doi.org/10.1016/S0893-133X(97)00194-2.

Nagata, E., A. de Toledo, and T. Oho. Invasion of Human Aortic Endothelial Cells by Oral Viridans Group Streptococci and Induction of Inflammatory Cytokine Production. Molecular Oral Microbiology 26, no. 1 (2011):78–88. doi: 10.1111/j.2041-1014.2010.00597.x.

Ni, Josephine, Ting-Chin David Shen, Eric Z. Chen, Kyle Bittinger, Aubrey Bailey, Manuela Roggiani, Alexandra Sirota-Madi, et al. “A Role for Bacterial Urease in Gut Dysbiosis and Crohn’s Disease.” Science Translational Medicine 9, no. 416 (November 15, 2017). https://doi.org/10.1126/scitranslmed.aah6888.

Patro, R., Duggal, G., Love, M. et al. Salmon provides fast and bias-aware quantification of transcript expression. Nat. Methods, (2017); 14: 417–419. doi: 10.1038/nmeth.4197

Pauwels, Lisa, Celine Maes, and Stephan P. Swinnen. “Aging, Inhibition and GABA.” Aging (Albany NY) 10, no. 12 (December 5, 2018): 3645–46. https://doi.org/10.18632/aging.101696.

Pyrkov T.V., Slipensky K., Barg M., et al. Extracting biological age from biomedical data via deep learning: too much of a good thing?. Sci Rep. (2018); 8(1):5210.. doi: 10.1038/s41598-018-23534-9

Quinn, Kylie M., Annette Fox, Kim L. Harland, Brendan E. Russ, Jasmine Li, Thi H.O. Nguyen, Liyen Loh, et al. “Age-Related Decline in Primary CD8+ T Cell Responses Is Associated with the Development of Senescence in Virtual Memory CD8+ T Cells.” Cell Reports 23, no. 12 (June 2018): 3512–24. https://doi.org/10.1016/j.celrep.2018.05.057.

Sae-Lee C., Corsi S., Barrow T.M., et al. Dietary Intervention Modifies DNA Methylation Age Assessed by the Epigenetic Clock. Mol Nutr Food Res. (2018); 62(23):e1800092. doi: 10.1002/mnfr.201800092

Toma, R., Duval, N., Pelle, N., Parks, M.M., Gopu, V., Torres, P.J., Camacho, F.R., Tily, H., Hatch, A., Perlina, A., Banavar, G., and Vuyisich, M.. A clinically-validated human capillary blood transcriptome test for global systems biology studies. Manuscript under review. doi: 10.1101/2020.05.22.110080

Wang, M., & Lemos, B. Ribosomal DNA harbors an evolutionarily conserved clock of biological aging. Genome Res. (2019); doi: 10.1101/gr.241745.118

Watson, M.M & Søreide, K. The Gut Microbiota influence on human epigenetics, health, and disease. Handbook of Epigenetics (2nd ed.) (2017); Academic Press. chap. 32, 495–510.

Yatsunenko T., Rey F.E., Manary M.J., Trehan I., Dominguez-Bello M.G., Contreras M., Magris M., Hidalgo G., Baldassano R.N., Anokhin A.P., Heath A.C., Warner B., Reeder J., Kuczynski J., Caporaso J.G., Lozupone C.A., Lauber C., Clemente J.C., Knights D., Knight R., Gordon J.I.. 2012. Human gut microbiome viewed across age and geography. Nature 486:222–227. doi: 10.1038/nature11053

## Supplementary References

Biagi, Elena, Claudio Franceschi, Simone Rampelli, Marco Severgnini, Rita Ostan, Silvia Turroni, Clarissa Consolandi, et al. “Gut Microbiota and Extreme Longevity.” Current Biology: CB 26, no. 11 (06 2016): 1480–85. https://doi.org/10.1016/j.cub.2016.04.016.

Cartron, Michaël L., Sarah Maddocks, Paul Gillingham, C. Jeremy Craven, and Simon C. Andrews. “Feo--Transport of Ferrous Iron into Bacteria.” Biometals: An International Journal on the Role of Metal Ions in Biology, Biochemistry, and Medicine 19, no. 2 (April 2006): 143–57. https://doi.org/10.1007/s10534-006-0003-2.

Claesson, Marcus J., Ian B. Jeffery, Susana Conde, Susan E. Power, Eibhlís M. O’Connor, Siobhán Cusack, Hugh M. B. Harris, et al. “Gut Microbiota Composition Correlates with Diet and Health in the Elderly.” Nature 488, no. 7410 (August 9, 2012): 178–84. https://doi.org/10.1038/nature11319.

Franceschi, C., M. Bonafè, S. Valensin, F. Olivieri, M. De Luca, E. Ottaviani, and G. De Benedictis. “Inflamm-Aging. An Evolutionary Perspective on Immunosenescence.” Annals of the New York Academy of Sciences 908 (June 2000): 244–54. https://doi.org/10.1111/j.1749-6632.2000.tb06651.x.

Gargari, Giorgio, Valentina Taverniti, Claudio Gardana, Cesare Cremon, Filippo Canducci, Isabella Pagano, Maria Raffaella Barbaro, et al. “Fecal Clostridiales Distribution and Short-Chain Fatty Acids Reflect Bowel Habits in Irritable Bowel Syndrome.” Environmental Microbiology 20, no. 9 (2018): 3201–13. https://doi.org/10.1111/1462-2920.14271.

Maraki, Sofia, and Ioannis S. Papadakis. “Rothia Mucilaginosa Pneumonia: A Literature Review.” Infectious Diseases 47, no. 3 (March 4, 2015): 125–29. https://doi.org/10.3109/00365548.2014.980843.

Mohan, P. K., L. V. Reddy, N. Satyanarayana, and K. Indira. “Age-Related Changes in Muscle Ammonia Detoxification Potential in Exhausted Rats.” Archives Internationales De Physiologie Et De Biochimie 95, no. 1 (March 1987): 37–42. https://doi.org/10.3109/13813458709075023.

Nagpal, Ravinder, Rabina Mainali, Shokouh Ahmadi, Shaohua Wang, Ria Singh, Kylie Kavanagh, Dalane W. Kitzman, Almagul Kushugulova, Francesco Marotta, and Hariom Yadav. “Gut Microbiome and Aging: Physiological and Mechanistic Insights.” Nutrition and Healthy Aging 4, no. 4 (n.d.): 267–85. https://doi.org/10.3233/NHA-170030.

Ravel, Jacques, Pawel Gajer, Zaid Abdo, G. Maria Schneider, Sara S. K. Koenig, Stacey L. McCulle, Shara Karlebach, et al. “Vaginal Microbiome of Reproductive-Age Women.” Proceedings of the National Academy of Sciences 108, no. Supplement 1 (March 15, 2011): 4680–87. https://doi.org/10.1073/pnas.1002611107.

Staerck, Cindy, Amandine Gastebois, Patrick Vandeputte, Alphonse Calenda, Gérald Larcher, Louiza Gillmann, Nicolas Papon, Jean-Philippe Bouchara, and Maxime J. J. Fleury. “Microbial Antioxidant Defense Enzymes.” Microbial Pathogenesis 110 (September 2017): 56–65. https://doi.org/10.1016/j.micpath.2017.06.015.

Volkov, Anton, Alena Liavonchanka, Olga Kamneva, Tomas Fiedler, Cornelia Goebel, Bernd Kreikemeyer, and Ivo Feussner. “Myosin Cross-Reactive Antigen of Streptococcus Pyogenes M49 Encodes a Fatty Acid Double Bond Hydratase That Plays a Role in Oleic Acid Detoxification and Bacterial Virulence.” Journal of Biological Chemistry 285, no. 14 (April 2, 2010): 10353–61. https://doi.org/10.1074/jbc.M109.081851.

Yoshikawa, Thomas T. “Antimicrobial Resistance and Aging: Beginning of the End of the Antibiotic Era?” Journal of the American Geriatrics Society 50, no. 7 Suppl (July 2002): S226–229. https://doi.org/10.1046/j.1532-5415.50.7s.2.x.

Zapata, Heidi J., and Vincent J. Quagliarello. “The Microbiota and Microbiome in Aging: Potential Implications in Health and Age-Related Diseases.” Journal of the American Geriatrics Society 63, no. 4 (April 2015): 776–81. https://doi.org/10.1111/jgs.13310.

Zhu, Feng, Yanmei Ju, Wei Wang, Qi Wang, Ruijin Guo, Qingyan Ma, Qiang Sun, et al. “Metagenome-Wide Association of Gut Microbiome Features for Schizophrenia.” Nature Communications 11, no. 1 (31 2020): 1612. https://doi.org/10.1038/s41467-020-15457-9.

Dinarello, C. A. “Interleukin-18.” Methods (San Diego, Calif.) 19, no. 1 (September 1999): 121–32. https://doi.org/10.1006/meth.1999.0837.

Sun, Xiaoting, and Qi Li. “Prostaglandin EP2 Receptor: Novel Therapeutic Target for Human Cancers (Review).” International Journal of Molecular Medicine 42, no. 3 (September 1, 2018): 1203–14. https://doi.org/10.3892/ijmm.2018.3744.

Manieri, Nicholas A., Eugene Y. Chiang, and Jane L. Grogan. “TIGIT: A Key Inhibitor of the Cancer Immunity Cycle.” Trends in Immunology 38, no. 1 (January 1, 2017): 20–28. https://doi.org/10.1016/j.it.2016.10.002.

Vazquez-Cintron, Edwin J., Ngozi R. Monu, Jeremy C. Burns, Roy Blum, Gregory Chen, Peter Lopez, Jennifer Ma, Sasa Radoja, and Alan B. Frey. “Protocadherin-18 Is a Novel Differentiation Marker and an Inhibitory Signaling Receptor for CD8+ Effector Memory T Cells.” PLOS ONE 7, no. 5 (May 2, 2012): e36101. https://doi.org/10.1371/journal.pone.0036101.

Monti-Rocha, Renata, Allysson Cramer, Paulo Gaio Leite, Maísa Mota Antunes, Rafaela Vaz Sousa Pereira, Andréia Barroso, Celso M. Queiroz-Junior, et al. “SOCS2 Is Critical for the Balancing of Immune Response and Oxidate Stress Protecting Against Acetaminophen-Induced Acute Liver Injury.” Frontiers in Immunology 9 (2019). https://doi.org/10.3389/fimmu.2018.03134.

Pal, Sangita, and Jessica K. Tyler. “Epigenetics and Aging.” Science Advances 2, no. 7 (2016): e1600584. https://doi.org/10.1126/sciadv.1600584.

Sun, Luyang, Ruofan Yu, and Weiwei Dang. “Chromatin Architectural Changes during Cellular Senescence and Aging.” Genes 9, no. 4 (April 16, 2018). https://doi.org/10.3390/genes9040211.

Pagiatakis, Christina, Elettra Musolino, Rosalba Gornati, Giovanni Bernardini, and Roberto Papait. “Epigenetics of Aging and Disease: A Brief Overview.” Aging Clinical and Experimental Research, December 6, 2019. https://doi.org/10.1007/s40520-019-01430-0.

Duron, Emmanuelle, Benoît Funalot, Nadège Brunel, Joel Coste, Laurent Quinquis, Cécile Viollet, Joel Belmin, et al. “Insulin-like Growth Factor-I and Insulin-like Growth Factor Binding Protein-3 in Alzheimer’s Disease.” The Journal of Clinical Endocrinology and Metabolism 97, no. 12 (December 2012): 4673–81. https://doi.org/10.1210/jc.2012-2063.

Bossù, Paola, Antonio Ciaramella, Maria Luisa Moro, Lorenza Bellincampi, Sergio Bernardini, Giorgio Federici, Alberto Trequattrini, et al. “Interleukin 18 Gene Polymorphisms Predict Risk and Outcome of Alzheimer’s Disease.” Journal of Neurology, Neurosurgery, and Psychiatry 78, no. 8 (August 2007): 807–11. https://doi.org/10.1136/jnnp.2006.103242.

Reitz, C., G. Tosto, B. Vardarajan, E. Rogaeva, M. Ghani, R. S. Rogers, C. Conrad, et al. “Independent and Epistatic Effects of Variants in VPS10-d Receptors on Alzheimer Disease Risk and Processing of the Amyloid Precursor Protein (APP).” Translational Psychiatry 3 (May 14, 2013): e256. https://doi.org/10.1038/tp.2013.13.

Lyons, Gretchen E., Tamson Moore, Natasha Brasic, Mingli Li, Jeffrey J. Roszkowski, and Michael I. Nishimura. “Influence of Human CD8 on Antigen Recognition by T-Cell Receptor-Transduced Cells.” Cancer Research 66, no. 23 (December 1, 2006): 11455–61. https://doi.org/10.1158/0008-5472.CAN-06-2379.

Zhang, Nu, and Michael J. Bevan. “CD8+ T Cells: Foot Soldiers of the Immune System.” Immunity 35, no. 2 (August 26, 2011): 161–68. https://doi.org/10.1016/j.immuni.2011.07.010.

